# Development of a lipid-based delivery system for the chemotherapeutic compound SN-38

**DOI:** 10.1101/792317

**Authors:** Alicia Soler Cantón, Niels van den Broek, Christophe Danelon

## Abstract

SN-38 is a chemotherapeutic compound with potent antitumor effects. However, its clinical application is currently limited due to its poor solubility and low stability at physiological pH. Liposomes and cyclodextrins have been long studied for the solubilization and delivery of hydrophobic compounds. Aiming to combine the advantages from both systems, we attempted to develop an SN-38-in-cyclodextrin-in-liposome formulation. We found that the encapsulation of SN-38-SBE-β-CD inclusion complexes in the lumen of liposomes was not possible, owing to the disassembly of liposomes and the formation of lipid nanoparticles, as revealed by size exclusion chromatography and single nanoparticle fluorescence microscopy. Interestingly, the retention time of SN-38 inside SN-38-SBE-β-CD-lipid nanoparticles is higher than in liposomes, whereby SN-38 was directly loaded into the lipid film. The toxicity of purified SN-38-SBE-β-CD-lipid nanoparticles was assayed in cultured cancer cells, showing no therapeutic advantage compared to bulk SN-38-SBE-β-CD complexes. Further formulation optimization, in particular an increased concentration of the nanoparticles, will be necessary to obtain cytotoxicity effects. Moreover, the results highlight the value of fluorescence imaging of single, surface-immobilized nanoparticles, in the development of liposomal delivery systems such as drug-in-cyclodextrin- in-liposomes.

## INTRODUCTION

SN-38, 7-ethyl-10-hydroxycamptothecin, is the active metabolite of Irinotecan (CPT-11), a compound approved by the Food and Drug Administration agency for use in the treatment of recurrent metastatic colorectal cancer. SN-38 interfers with the human topoisomerase I, impeding both DNA replication and transcription [1] [2].

The therapeutic activity of CPT-11 depends on its conversion efficiency to SN-38. Upon administration, only 2 to 8% of CPT-11 is metabolized into the active SN-38 through carboxylesterase mediated cleavage in the liver [3, 4]. The conversion rate is relatively dependent on genetic variability, making it unpredictable, posing significant life-threatening risks and complicating clinical management of patients [3, 5]. Unlike its precursor, SN-38 does not require activation in the liver. Moreover, *in vitro* toxicity studies indicate that SN-38 is up to 1,000-fold more potent than CPT-11 against different tumor cell lines [5]. Additionally, the biological half-life of SN-38 is much longer than that of CPT-11 [3]. Thus, SN-38 holds great pharmaceutical advantage over CPT-11 as an anticancer candidate.

Despite its potent activity toward tumor cells *in vitro*, clinical application of SN-38 remains challenging due to its extremely poor solubility in aqueous solutions and other pharmaceutically acceptable solvents [6]. SN-38 exists in two pH-dependent reversible forms: an active lactone ring at acidic pH and an inactive carboxylate form at basic pH (Fig. 1a). The closed lactone moiety is a required feature for the anti-tumor activity of SN-38. Unfortunately, at physiological pH, the active lactone is partially converted into the inactive carboxylate that becomes dominant at equilibrium [3]. In addition, *in vivo*, the carboxylate form tightly binds to human serum albumin with an affinity 150-fold higher than the lactone form, further shifting the lactone-carboxylate equilibrium towards the inactive state [7]. Therefore, the instability of the lactone SN-38 under physiological conditions represents a major challenge towards an effective therapy with SN-38.

**Figure 1.**
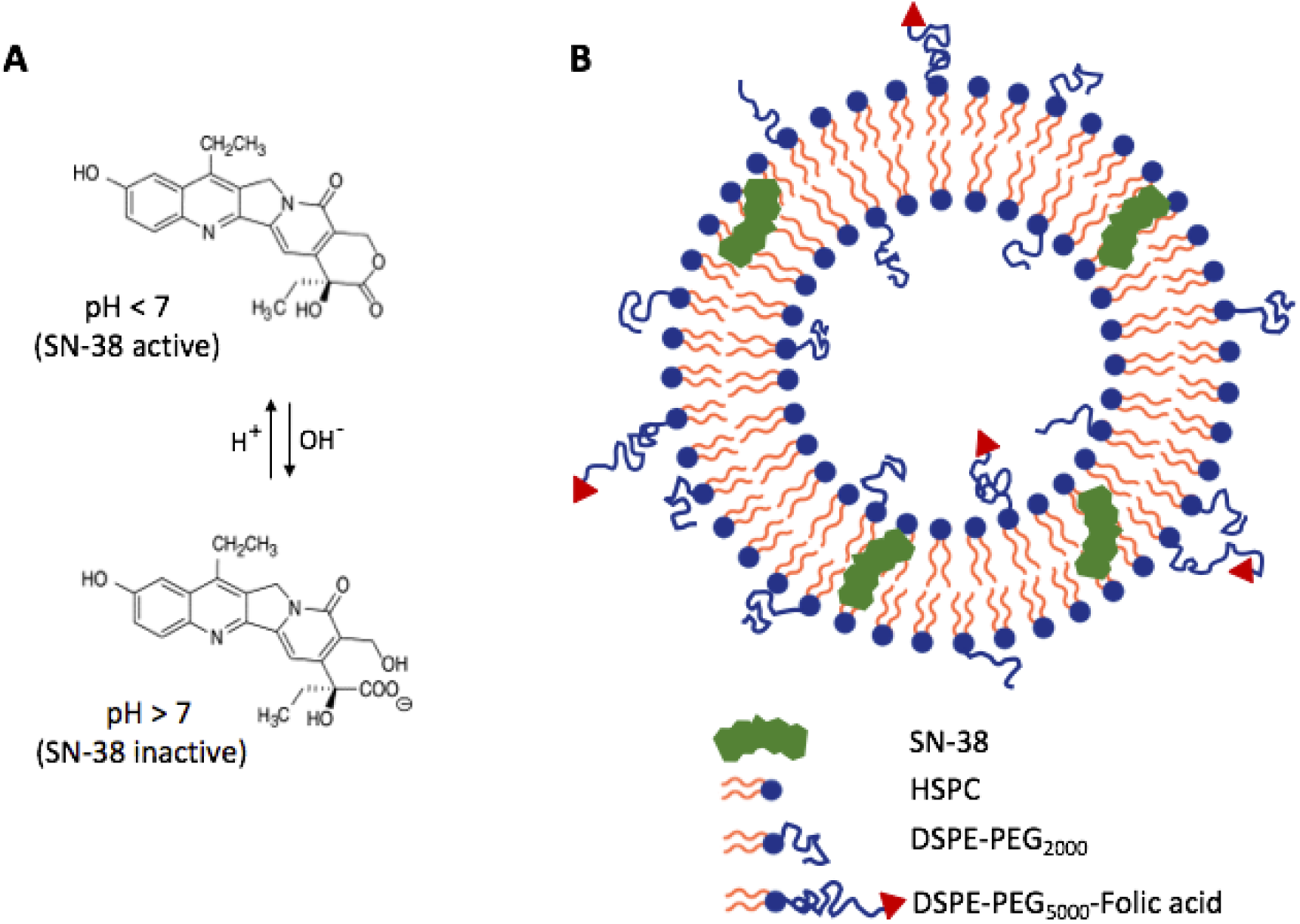
A) Chemical structure of the lactone and carboxylate forms of SN-38. Adapted from [4]; B) Schematic of SN-38-loaded folate-targeted liposomes.

In this work, we aimed at developing two types of lipid-based drug delivery systems for SN-38 in order to overcome some of the above challenges: SN-38-loaded liposomes and SN-38-in- cyclodextrin-in liposomes. Liposomal delivery systems can solubilize hydrophobic drugs in their lipid bilayer [8], improve their stability [9] and tumor targeting ability *in vivo* [10], and reduce their systemic toxicity. Previous efforts have attempted to incorporate camptothecin derivates into liposomes [5]. It has been shown that interaction with a lipid bilayer stabilizes the lactone ring moiety of camptothecin drugs by protecting it from hydrolysis [11]. Furthermore, liposomes prevent the complexation of the carboxylate form with human serum albumin [12]. In 2004, Zhang et al. [3] described the development of a non-targeted SN-38 liposomal formulation (LE- SN-38) that improved the pharmacokinetic profile, safety and tolerability of SN-38 in its phase I trial investigation for the treatment of metastatic colorectal cancer [13].

Although liposomalization of drugs can enhance their tumor distribution passively through the enhanced permeability and retention effect [14–16], insufficient uptake at tumor sites and nonspecific association to healthy tissues decrease their therapeutic efficacy and may provoke harmful side effects [17]. Active targeted delivery increases tissue selectivity and thus reduces side effects [18]. Folic acid has been extensively investigated as a ligand for tumor-specific and targeted drug delivery because its receptors (folic acid receptors, FR) are overexpressed on the surface of a variety of malignant tumor cells, while being minimally distributed in healthy tissues [19]. Folate-conjugated lipidic nanocarriers have been shown to increase drug accumulation in FR-overexpressing tumors [20]. Different chemotherapeutic agents have been delivered using liposomes conjugated to the folate ligand via a polyethyleneglycol spacer [21]. Recently, a study has shown that SN-38-loaded, folate-functionalized liposomes can target breast cancer tumors and reduce unspecific SN-38 toxicity *in vivo* [22].

Incorporation of hydrophobic drugs in the lipid bilayer of liposomes is always restricted in terms of the drug-to-lipid mass ratio [23, 24]. Thus, new strategies to further enhance loading efficiency of highly lipophilic camptothecin compounds, like SN-38, into liposomes must be deployed. Cyclodextrins are naturally existing water-soluble cyclic oligosaccharides that form inclusion complexes by sequestering small molecules or moieties of larger molecules inside their hydrophobic cavity [25, 26]. Drugs complexed with cyclodextrins show increased solubility [27], stability [28], dissolution rate and bioavailability [29], leading to enhanced cytotoxicity of several anti-cancer drugs, such as camptothecin, curcumin, doxorubicin, docetaxel and paclitaxel [28]. Sulfobutyl ether-β-cyclodextrin (SBE-β-CD, marketed as Captisol™) is used in various drug formulations. Recently, Vangara et al. [28] found that SN-38 could form stable inclusion complexes with SBE-β-CD, enhancing its solubility by more than 1,000 fold and improving its anticancer potency on ovarian cells. However, the relatively poor affinity of the drug- cyclodextrin complex leads to rapid dissociation and premature clearance of the drug after intravenous injection [30–34].

The entrapment of drug-cyclodextrin complexes in the aqueous phase of liposomes might circumvent the problems associated to the conventional integration of drugs into the lipid phase of liposomes or into the hydrophobic cavity of cyclodextrins. Drug-in-cyclodextrin-in-liposomes could provide an ideal integrative system combining the assets of both approaches [33–35]. In this study, we examined the potential of two different lipid-based systems for tumor-targeted delivery of SN-38. First, we prepared a formulation of folate-labeled, SN-38-loaded liposomes (Fig. 1b). To achieve long-term storage of liposome precursors and facilitate laboratory distribution, we prepared SN-38-loaded lipid films coated onto glass beads. Furthermore, we prepared a folate-labeled SN-38-in-cyclodextrin-in-liposome formulation. Particle size distribution, drug retention time and association/internalization with/by human carcinoma FR- overexpressing KB cells were investigated.

## MATERIALS AND METHODS

### Materials

SN-38 was purchased from Seleckchem (Houston, TX, USA), diluted in DMSO to 50 mM (20 mg/ml), aliquoted and stored at –80 °C. Captisol™ was kindly provided by CyDex Inc. Pharmaceuticals (USA). HeLa cells were kindly provided by Eric P. van der Veer, Leiden University Medical Center, Leiden (The Netherlands). KB cells (ACC 136), a derivative cell line from HeLa, were purchased from DSMZ. Hydrogenated L-α-phosphatidylcholine (HSPC, Avanti Polar Lipids) powder was kindly provided by to-BBB Inc (Leiden). 1,2-distearoyl-*sn*-glycero-3- phosphoethanolamine-N-[biotinyl(polyethylene glycol)-2000] (DSPE-PEG-biotin) 10 mg/ml, cholesterol 50 mg/ml and 1,2-distearoyl-*sn*-glycero-3-phosphoethanolamine-N- [folate(polyethylene glycol)-5000] (DSPE-PEG-folate) 500 µg were purchased from Avanti Polar lipids. Texas Red^®^ 1,2-dihexadecanoyl-*sn*-glycero-3-phosphoethanolamine (DHPE-TR) was purchased from Invitrogen.

### Formation of SN-38-containing lipid films

A 50 mM stock solution of SN-38 in DMSO was diluted in methanol at a final concentration of 10 mM. The sample was further diluted ten times in chloroform and 50 µl were added to a lipid solution composed of 30 µl of HSPC 10 mg/ml (stock concentration), 2 µl of cholesterol 50 mg/ml, 10 µl of DSPE-PEG-biotin 10 mg/ml and 5 µl of DHPE-TR 1 mg/ml, all dissolved in chloroform, resulting in a molar ratio HSPC:cholesterol:DSPE-PEG-biotin:DHPE-TR of 56.7:37.8:4.8:0.5 and a total mass of lipids of 0.5 mg. A negative control sample devoid of SN-38 was prepared by mixing 1 µl of DMSO and 49 µl methanol, and by proceeding as described above. The resulting samples were added to 212-300-µm glass beads (Sigma Aldrich) in a round-bottom glass flask at a lipid-to-bead mass ratio of 1:760. The solvent was rotary evaporated at room temperature and 500 mbar overnight and the pressure was gradually decreased down to 20 mbar. Then, temperature was increased to 80 °C and rotary evaporation was continued for 3 h at 20 mbar to remove traces of DMSO. Lipid film-coated beads were collected in 2-ml amber glass vials (Sulpeco) and flushed with argon for a few seconds before storage. Vials were closed, sealed with Parafilm and stored at –20 °C until use.

### Formation of folate-containing lipid films

A lipid mixture in chloroform with a molar ratio of HSPC:cholesterol:DSPE-PEG-biotin:DHPE- TR:DSPE-PEG-folate equal to 56.6:37.8:4.8:0.5:0.1 for a total mass of 1 mg was added to 212- 300-µm glass beads in a round-bottom glass flask at a lipid-to-bead mass ratio of 0.003:1. The solvent was rotary evaporated at about 40 rpm and 200 mbar overnight and 20 mbar for 10 min. Lipid-coated beads were stored at –20 °C and were used within two weeks.

### Liposomal SN-38 production

Lipid-SN-38-coated beads were desiccated for 20 min at room temperature before use. About 50 mg of functionalized beads were poured in a reaction tube and immersed in 200 µl of swelling solution consisting of acidic PBS (pH 5.3). Liposome production was induced by vigorous vortex for 1 min, leading to a final lipid concentration of 0.34 mg/ml and a final SN-38 concentration of 33.3 µM. SN-38-containing liposomes were extruded in a mini-extruder (Avanti Polar Lipids) using a 0.2 µm pore-size polycarbonate membrane (Avanti Polar Lipids) and two 10 mm filter supports (Whatmann) by 11 to 13 extrusion passages. The obtained small unilamellar vesicles (SUVs) were incubated at 4 °C for 48 or 72 h protected from light.

### Preparation of folate-labeled liposomes

Approximately 50 mg of folate-labeled lipid-coated beads were briefly flushed with argon and immersed in 200 µl of PBS. Liposome production was induced by natural swelling for 2 h with manual rotation every 15 min at 60 °C (HSPC *T*_m_ = 53 °C). The final lipid concentration was 0.8 mg/ml. Liposomes were downsized by extrusion to obtain SUVs as described above, except that temperature was set to 60 °C.

### Dynamic light scattering

The size distribution of liposomes was analyzed by dynamic light scattering (DLS). For a typical experiment, 10 µl of the SUV solution was diluted in 90 µl of PBS buffer and placed into a disposable microcuvette (ZEN0040, Malvern). The measurements were carried out on a Zetasizer Nano ZS (Malvern, UK) operating at a scattering angle of 173° at room temperature. For each condition, each run consisted of three measurements of 12-20 acquisitions per sample. Liposome colloidal stability was studied over time by monitoring their mean size and size distribution every day for 5 days upon storage at 4 °C.

### Size exclusion chromatography

A homemade size exclusion chromatography column was prepared by filling the neck of a 150 mm Pasteur glass pipette with cotton. The cotton was washed with MilliQ water and 1.7 ml or 2 ml of Sepharose beads 4B (45-165 µm bead diameter) were added. Columns were packed overnight at room temperature. Prior to usage, columns were washed three times with a volume equal to that of the beads added. Column washes were done with either sample buffer or ammonium acetate (25 mM, pH 8.6) depending on the experiment. Liposome samples pre-incubated for 0, 48 or 72 h at 4 °C were added when the eluent level reached the top of the gel. Buffer flow was driven by gravity only. The sample was allowed to penetrate the gel entirely prior to adding repetitively small volumes of buffer for continuous elution. A sample volume between 30 and 150 µl was purified in each column, and 30-48 elution samples were collected in microcentrifuge tubes. Two drops of eluate were collected per tube.

### Fluorescence spectroscopy

Elution samples of 10 µl were dispensed in wells of a black 384-well microplate (Greiner Bio One). Fluorescence intensity was measured with a CLARIOstar® Microplate reader (BMG LABTECH’s) using 379/10 nm excitation and 548/10 nm emission wavelengths for SN-38, and 562/15 nm excitation and 606/10 nm emission wavelengths for the liposome membrane dye (Texas Red). A focal height of 11 mm was set for all measurements. Gains of 1000, 1500, 2000, or 3000 for SN-38 and Texas Red were used depending on the experiment. The emission spectra of SN-38 and Texas Red were measured on a glass-bottom 384-well microplate (µClear®, non-binding, Greiner Bio One) using excitation wavelengths of 379/10 nm and 562/10 nm, respectively, a gain of 3000 and a focal height of 3 mm.

### Preparation of chambered coverslips

Chambered coverslips were fabricated to image liposomes. First, two holes with diameters of 5 and 7 mm were drilled into a coverslide (76 × 26 mm, Menzel-Gläser) with a Sand Blaster. After a brief washing step with MilliQ water, coverslides were consecutively sonicated in a bath (Branson 1510 ultrasonic cleaner, Ultrasonics) with 2% Hellmanex and MilliQ water for 10 min and with 100% ethanol. Dried coverslides were plasma cleaned in a Harrick Plasma PDC-002. Coverslips (24 × 60 mm, Menzel-Gläser) were also cleaned following the protocol described above. A drop of Norland optical adhesive (NOA 81, Ultraviolet curing) was applied to the surroundings of the apertures on a single side of the coverslides, and a clean coverslip was immediately sealed on top by 5 min irradiation with a 365-nm light using a Promed UVL 36 lamp. The chambered coverslips were incubated for 48 h at 55 °C, washed as mentioned above, and stored in clean Petri dishes until usage.

### Fluorescence confocal microscopy

The 5-mm wide well of the chambered coverslip was functionalized with 10 µl of 1 mg/ml BSA-Biotin (Thermo Fisher Scientific) for 5 min, washed three times with MilliQ water, incubated with 10 µl of 1 mg/ml Neutravidin (Thermo Fisher Scientific) for 5 min at room temperature, washed three times with MilliQ water and finally incubated with 10 µl PBS. Before imaging, PBS was replaced with 7-10 µl of the lipid sample (liposomes or nanoparticles). The 7-mm hole was filled with 15 µl MilliQ water to avoid sample evaporation during imaging. After sample addition, chambered coverslips were sealed with a 1-mm thick silicone spacer (Press-to-seal silicone sheet, Life technologies) and a clean coverslip. Imaging was performed using a laser scanning confocal microscope (A1+, Nikon) equipped with a ×100 oil immersion objective. The 405 nm and 561 nm laser lines with appropriate dichroic mirrors and emission filters were used to image SN-38 and Texas Red, respectively.

### High performance liquid chromatography

SN-38 detection was performed using a reverse-phase HPLC system (Agilent 1260 Infinity Quaternary LC, Santa Clara, United States) consisting of a quaternary pump, an autosampler module and a fluorescence detector set at 368 nm and 515 nm for excitation and emission wavelengths, respectively. The mobile phase consisted of freshly prepared 25 mM ammonium acetate pH 5.5 (solvent A) and acetonitrile (solvent B) and the column was an Agilent SB-C18 (50 mm × 2.1 mm, 1.8 µm). The mobile phase was pumped at a flow rate of 0.5 ml/min in accordance with the following program (time in minute, percent solvent B): 0, 15; 10, 40; 10.01, 100; 11, 100; and 11.01, 15. The total run time was 15 min. The elution times of the SN-38 lactone and carboxylate forms were 6.05 and 1.75 min, respectively. When high amounts of SN-38 were injected, the washing step of the column was extended by one minute and the injection needle was washed with 1:1 (v:v) methanol/water after each injection. Additionally, 10 µl of acetonitrile were injected between sample runs to clean the column and avoid sample-to-sample contaminations. The flow rate was 0.5 ml/min and the maximum pressure in the column was 600 bar. Samples were loaded into low-volume HPLC vial inserts (250 µl, Agilent) in 2-ml screw top vials (Agilent) and kept at 4 °C until injection. All samples containing liposomes or lipid nanoparticles were pre-diluted ten times in methanol, vortexed for 30 sec and spun down in a tabletop centrifuge for 30-60 sec to remove any precipitates. HPLC samples containing 90% methanol were injected soon after preparation to avoid conversion of carboxylate SN-38 into lactone SN-38. Sample injection volumes were 1 µl, 2 µl or 10 µl depending on the experiment.

### pH dependence assays

The 50 mM SN-38 stock solution in DMSO was diluted hundred times in acidic PBS (pH 5.3). This sample was further diluted fifty times in either acidic PBS, neutral PBS (pH 7.4) or in 50 mM sodium phosphate (pH 8.3) leading to a final SN-38 concentration of 10 µM. Tubes were then sealed, shielded from light and incubated at 37 °C overnight. Before HPLC measurements, samples were spun for 20 min at 13,200 rpm, and 10 µl were carefully harvested and injected into the HPLC.

For the SN-38 inactivation kinetics study, the SN-38 stock solution was diluted in acidic PBS at 32.5 µM final concentration. This sample was further diluted twenty times in either acidic PBS or in 50 mM sodium phosphate (pH 8.3), and was incubated at 37 °C in sealed tubes. A 20 µl aliquot was harvested from each tube at different time points: 1 h, 2 h, 3 h, 4 h, 5 h and 24 h. All samples were diluted ten times in methanol giving a final SN-38 concentration of 1.7 µM, vortexed for 30 sec, spun for 20 min at 13,200 rpm and 10 µl were injected into the HPLC.

### SN-38 release experiments

A 150-µl sample of SN-38-containing SUVs was incubated at 4 °C for 48 h and purified by size exclusion chromatography through a 1.7-ml column pre-washed with acidic PBS. A total of 48 elution samples were collected in microcentrifuge tubes. Fluorescence intensity of SN-38 and Texas Red in each sample was analyzed on a CLARIOstar^®^ spectrofluorometer. The samples containing the highest membrane dye signal (liposome population) were pooled. A 20-µl aliquot was used for HPLC analysis (time zero), while the remaining volume (195 µl) was re-purified through a new size exclusion chromatography column with a 20-min lagtime after the first purification. After measuring the SN-38 and Texas Red fluorescence signals from all eluates of the second purification, the fractions corresponding to the liposome population were pooled and 20 µl were used for a second HPLC inspection.

### Liposome-mediated SN-38 stability assays

A 10-µl sample of SN-38-containing SUVs was incubated at 4 °C for 72 h, transferred in a new tube and diluted twenty times in ammonium acetate (25 mM, pH 8.6), resulting in 1.6 µM SN- 38 final concentration. A 20-µl aliquot was used for HPLC analysis (unpurified liposomes, time zero). The remaining solution was incubated for 2 h at 37 °C in sealed tubes shielded from light and a 20-µl aliquot was harvested for HPLC analysis (unpurified liposomes, time 2 h). SN-38- containing SUVs were purified by size exclusion chromatography as described above and 20 µl of the pooled liposome-containing fractions were analyzed by HPLC (purified liposomes, time zero). The remaining solution was incubated for 2 h at 37 °C in sealed tubes shielded from light and a 20-µl aliquot was harvested for HPLC analysis (purified liposomes, time 2 h). Negative controls were conducted in the absence of lipids. A sample of 32.5 µM SN-38 in acidic PBS was prepared as described above and further diluted twenty times in ammonium acetate (25 mM, pH 8.6), resulting in 1.6 µM SN-38 final concentration. A 20-µl aliquot was harvested for HPLC analysis (SN-38 bulk, time zero). The remaining solution was incubated for 2 h at 37 °C in sealed tubes shielded from light and a 20-µl aliquot was harvested for HPLC analysis (SN-38 bulk, time 2 h).

### Complexation of SN-38 with SBE-β-CD

Solubility of SN-38 was studied in the presence of varying amounts of SBE-β-CD in 1 ml final volume of acidic PBS. First, a 200-mM SBE-β-CD solution was prepared by dissolving 0.215 g of SBE-β-CD in 500 µl of acidic PBS. Samples of 800 µM, 10 mM, 48 mM, 100 mM and 150 mM SBE- β-CD were prepared by serial dilutions in the same buffer. All tubes were vortexed and incubated at 37 °C until all SBE-β-CD was dissolved. Next, SN-38 was added to a final concentration of 100 µM in each tube, resulting in samples with the following SN-38 to SBE-β-CD molar ratios: 1:8, 1:100, 1:480, 1:1000 and 1:1500. Samples were sonicated for 15 min in a bath sonicator, tumbled overnight at room temperature to form SN-38-SBE-β-CD complexes, centrifuged for 5 min at 13,200 rpm, and 900 µl were carefully transferred to new tubes in order to remove undissolved precipitates. Quantitation of SN-38 solubilization was carried out by measuring SN- 38 fluorescence intensity by HPLC (1-µl sample injection) and by spectrofluorometry using a plate reader, as described above. For each technique, a solubility curve was generated by plotting the measured fluorescence intensity values (plate reader) or the chromatogram peak areas (HPLC) against the molar excess of cyclodextrin used in each condition.

We also prepared samples with lower ratios of SN-38 to SBE-β-CD, namely 1.25:1 and 1:8, and varying absolute amounts of both molecules. All samples were sonicated for 15 min and rotated at 20 rpm overnight at room temperature. Aliquots of 600 µl from each sample were filtered with a 0.45-µm filter to remove undissolved SN-38 and 2 µl were injected for HPLC analysis. Moreover, the protective effect of SBE-β-CD on the lactone SN-38 was investigated in samples containing 25 µM SN-38 and 10.2 mM SBE-β-CD (or none in negative controls, the equivalent volume was substituted with MilliQ water) in 50 mM sodium phosphate buffer (pH 8.3). Sample preparation was as described above for the solubility study. Preparation of SN-38-SBE-β-CD complexes for cell viability assays was performed as described above, except that final concentrations of 1 mM SN-38 and 8 mM SBE-β-CD in acidic PBS were used.

### Quantitation of SN-38

Ten standards consisting of 100, 50, 25, 5, 2, 1, 0.5, 0.1, 0.05 and 0.01 µM of SN-38 in acidic PBS supplemented with a molar excess of SBE-β-CD were prepared. A minimum molar ratio of SN- 38:SBE-β-CD of 1:1000 was used by adding 100 mM SBE-β-CD to ensure complete SN-38 solubilization. First, 200 mM SBE-β-CD stock solution was prepared in PBS by dissolving 1.72 g in 4 ml of acidic PBS. The tube was vortexed and incubated at 37 °C until SBE-β-CD was dissolved, and the appropriate volume was added to the ten SN-38 standard samples, each having a final volume of 500 µl. All samples were sonicated for 15 min in a bath sonicator, tumbled overnight at room temperature in sealed tubes protected from light, spun for 5 min at 13,200 rpm and 400 µl were carefully transferred to new tubes. SN-38 fluorescence signal was measured by spectrofluorometry and HPLC as described above. Two calibration curves were generated by plotting the measured SN-38 fluorescence intensity values and chromatogram peak areas for each standard against their predetermined amounts. The standard curves were then used to calculate the concentration of SN-38 added to cells in cytotoxicity assays.

### Preparation of SN-38-SBE-β-CD-lipid nanoparticles

Folate-labeled lipid-coated beads were briefly flushed with argon and about 50 mg were immersed in 200 µl of swelling solution consisting of either 8 mM SBE-β-CD or 1 mM SN-38 complexed with 8 mM SBE-β-CD in acidic PBS. Liposome production was carried out by natural swelling for 2 h at 60 °C with gentle manual rotation of the tube every 15 min. Final lipid concentration was 0.8 mg/ml in the two samples. Liposomes were extruded as described above, except that temperature was set to 60 °C. Right after extrusion, the resulting SUV solutions were directly purified by size exclusion chromatography through freshly prepared columns (see above). The column was washed six times with 1 ml neutral PBS (pH 7.12) prior addition of 125 µl of an SUV solution. A total of 48 eluates per sample were collected in microcentrifuge tubes. The SN-38 and membrane dye fluorescence was measured as previously described.

### Cell culture

HeLa and KB cells were cultured in folate-deficient RPMI-1640 medium (ThermoFisher) supplemented with 10% heat-inactivated fetal bovine serum (ThermoFisher) with 5% CO2 at 37 °C. Cells were discarded after 35 passages.

### Fluorescence imaging of KB cells

3 × 10^5^ KB cells were seeded on glass-bottom microscopy plates (MatTek; 9.5 cm^2^) and incubated for 24 h at 37 °C with 5% CO_2_ prior to imaging. Then, 40 µl of either the folate-decorated HSPC- liposomes or the SN-38-SBE-β-CD lipid nanoparticles were added to the monolayers of KB cells. To assay the selective binding of the folate-decorated HSPC-liposomes with KB cells, a control experiment was performed, in which the cell medium was supplemented with 5 mM soluble folic acid during incubation with the SUVs. After 2-h incubation, the microscopy plates were mounted on the stage of a confocal microscope (see above for details) equipped with an environment control chamber set to 37 °C and 5% CO_2_. When indicated, time-lapse imaging was carried out during 2 h.

### Cell viability assay

100-µl cell suspensions with a density of 10 cells/µl were seeded on a white flat-bottom opaque walled 96-well plate (Sigma Aldrich) and cultured in RPMI-1640 medium supplemented with 10% heat-inactivated fetal bovine serum and penicillin/streptomycin (ThermoFisher) at 37 °C and 5% CO_2_ overnight. A volume of 12 to 15 µl of purified SN-38-SBE-β-CD-lipid nanoparticles was added to KB and HeLa cells. In a series of controls, the SN-38-SBE-β-CD-lipid nanoparticles solution was substituted with either PBS pH 7.4, a bulk SN-38-SBE-β-CD solution with an SN-38 concentration adjusted to that of the elution pool of SN-38-SBE-β-CD, purified SBE-β-CD-lipid nanoparticles with a lipid concentration adjusted to that of the purified SN-38-SBE-β-CD-lipid nanoparticles, an SN-38-SBE-β-CD bulk solution with 1 mM SN-38 and 8 mM SBE-β-CD, or a solution with 8 mM SBE-β-CD. All conditions were assayed in triplicate. Cell viability was performed after 72 h incubation according to the CellTiter-Glo^®^ protocol (Promega). Luminescence was measured with a CLARIOstar^®^ Microplate Reader (BMG LABTECH’s) set to detect light with wavelength between 520 nm and 620 nm, and a focal height of 11 mm.

## RESULTS

### Liposome production by swelling of SN-38-loaded lipid films supported on glass beads

SN-38-containing liposomes were prepared by lipid film hydration. A lipid composition mimicking that of Doxil^®^ was supplemented with three types of functionalized phospholipids conjugated to either folate as a cell targeting agent, Texas Red as a fluorescent membrane reporter and to biotin through a polyethylene glycol linker for enhanced immobilization during imaging (Fig. 1b). To facilitate storage, handling and distribution of lipid mixtures, while increasing the solubilization of SN-38, we prepared lipid-coated glass beads with SN-38 incorporated in the lipid film. Upon rehydration, multilamellar vesicles are produced as precursors of 200-nm unilamellar vesicles prepared by extrusion. Formulation of lipid-SN-38- coated beads as precursors of SN-38-containing small unilamellar vesicles is particularly suited because large batches can be prepared at once and be easily distributed across laboratories in the form of solvent-free aliquots. The functionalized beads could be stored for more than 6 months at –20 °C without noticeable conversion of SN-38 from the lactone to the carboxylate form or liposome stability.

Liposomes were prepared in acidic PBS (pH 5.3) to maintain SN-38 in its active state. SN-38- doped HSPC-liposomes were immobilized on a glass coverslip and imaged by fluorescence microscopy. The SUVs were localized through their membrane dye. They appear as discrete diffraction-limited spots with a width <1 µm (Fig. 2a). The population-wide liposome size distribution was determined by light scattering. Both SN-38-loaded SUVs and control SUVs had a similar average diameter of 225 and 216 nm, respectively, 7 days after liposome production (Fig. 2b, Fig. S1). These results indicate that the inclusion of SN-38 does not influence vesicle size, aggregation state or stability.

**Figure 2.**
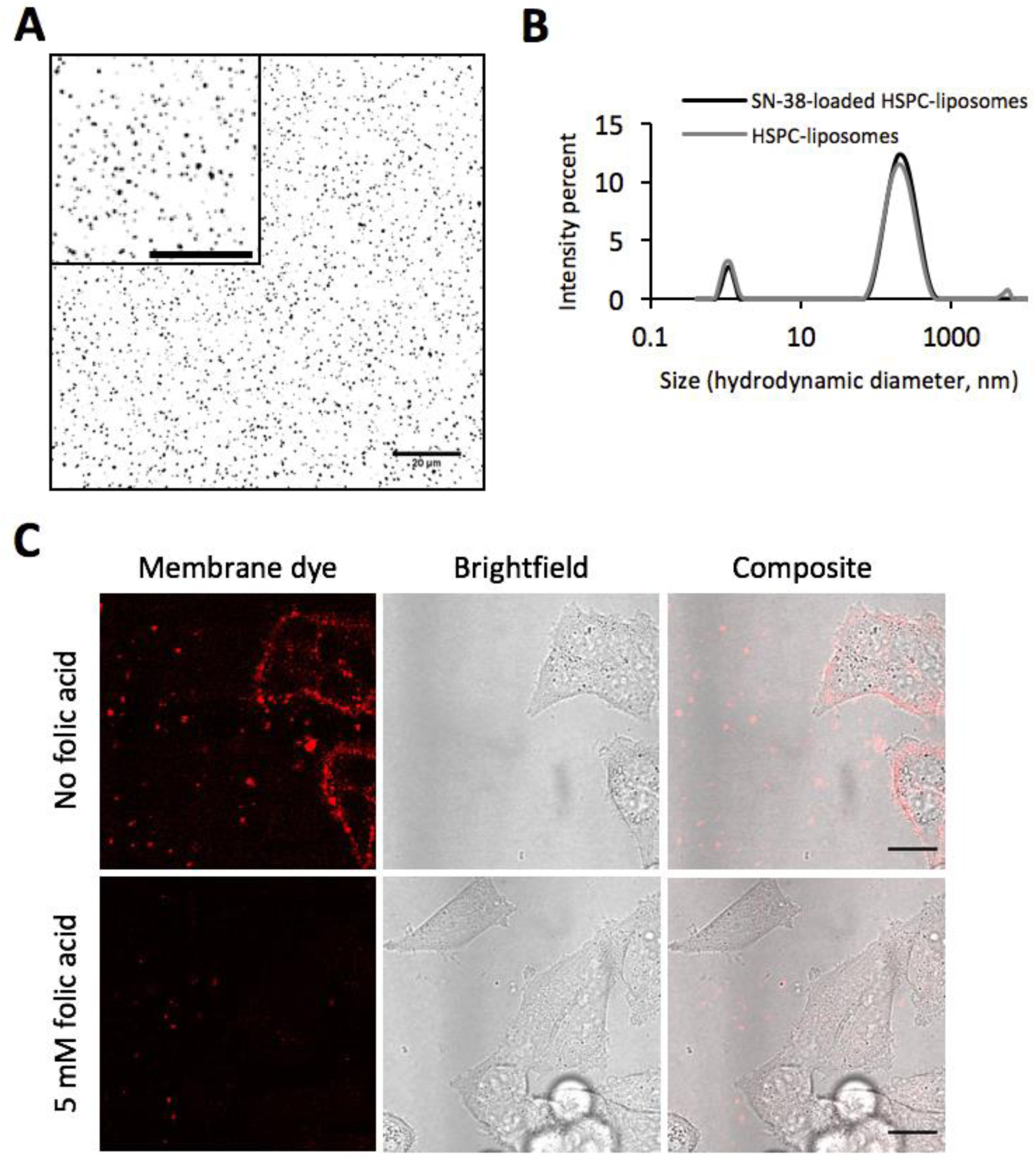
A) Fluorescence confocal images of SN-38-doped HSPC-liposomes tethered on a glass coverslip. The Texas Red membrane dye channel is shown. Single liposomes correspond to diffraction-limited spots. The inset is a zoom-in image from a larger field of view. Scale bars are 20 µm. B) Liposome size distribution as measured by dynamic light scattering. The SN-38-doped HSPC-SUV and HSPC-SUV samples were analyzed after 7 days at 4 °C. C) Fluorescence and brightfield confocal images of cultured KB cells in the presence of folate-labeled HSPC-liposomes incubated for 2 h at 37 °C. In a control sample, 5 mM folic acid was supplemented during incubation with the vesicles. Scale bars are 20 µm.

The ability of folate-modified HSPC-liposomes to target living cells was investigated in the absence of SN-38 using cultured KB cells. The specific role of the folate ligand in cellular interaction was evaluated by performing a competitive binding assay in which 5 mM of free folic acid was added to the medium to saturate the cell-surface folate receptors. Fluorescence imaging up to 2 h after incubating liposomes with the cells revealed that SUVs interact with the plasma membrane of KB cells mostly in the absence of free folate (Fig. 2c), demonstrating that folate ligand promotes liposome binding. Moreover, several vesicles clearly localized on the inner side of the plasma membrane (Fig. 2c) suggesting internalization during endocytosis of the folate receptors. These results prompted us to study the potential of folate-conjugated HSPC- liposomes for the targeted delivery of SN-38.

### SN-38 is rapidly released from folate-HSPC liposomes

The stability of SN-38 in the liposome membrane was investigated by size exclusion chromatography to quantify the amounts of free and liposome-loaded SN-38 molecules. The spectrally separated SN-38 and membrane dye fluorescence signals allowed us to monitor their respective levels in the course of elution. The corresponding intensity profiles are shown in Fig. 3a, b. Folate-modified HSPC-liposomes were identified as a single peak spanning across eight fractions, both in the presence and absence of SN-38. SN-38 mostly eluted as one peak after fraction 20, representing the free drug that has been released from the SUVs. Interestingly, a fraction of SN-38 co-eluted with liposomes when the sample was left to pre-incubate for 48 h prior column purification (Fig. 3b). Although the underlying mechanism requires clarification, we decided to quantify the rate of release under this condition and to determine the amounts of active (lactone) and inactive (carboxylate) SN-38.

**Figure 3.**
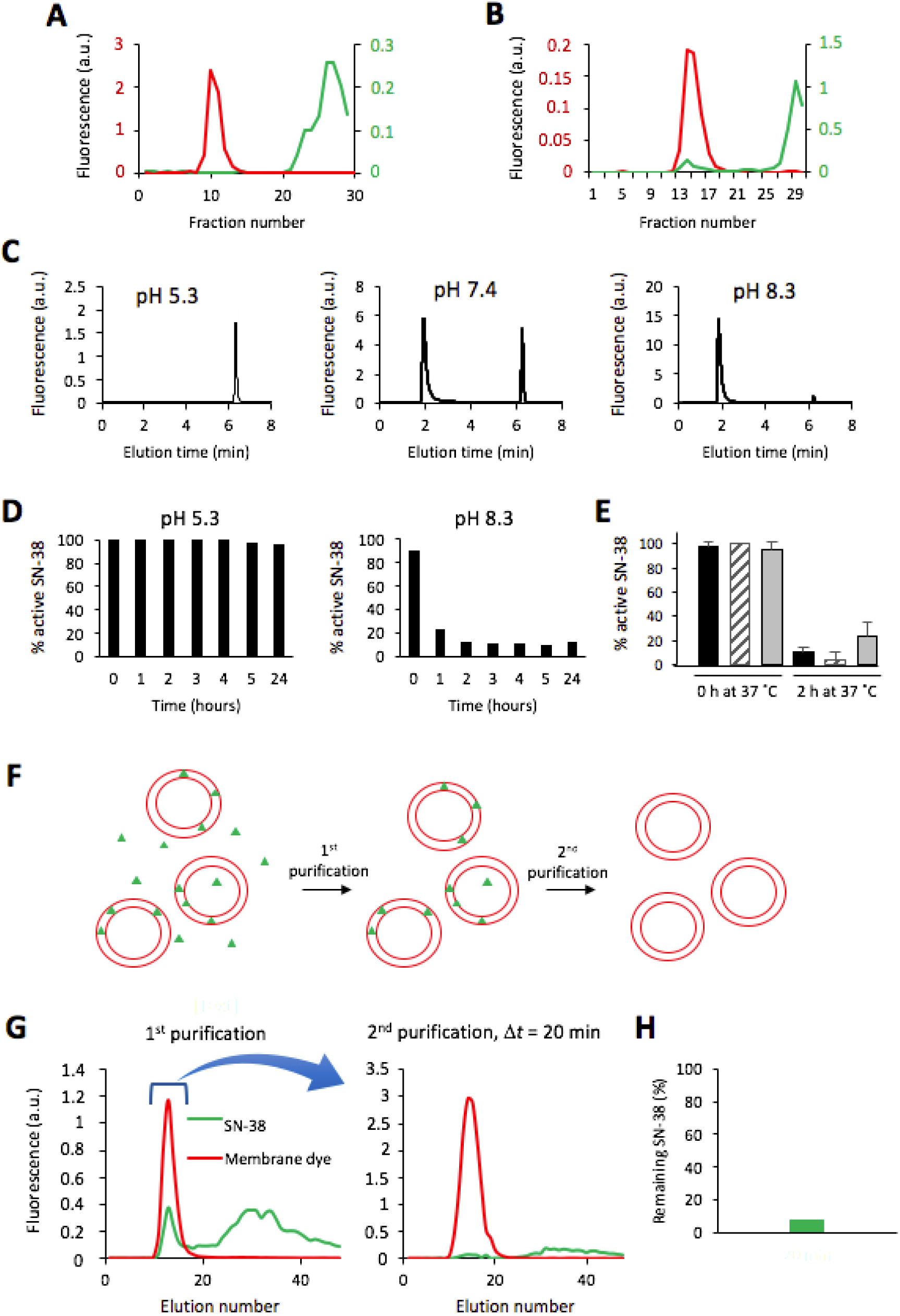
A) Elution profiles of SN-38 (green) and liposome (red) fluorescence intensity in size exclusion chromatography experiments with SN-38-loaded HSPC-liposomes. The sample was directly transferred to the purification column (A) or was incubated for 48 h at 4 °C prior to purification (B). C) HPLC chromatograms of SN-38 (10 µM) incubated overnight at 37 °C in acidic PBS (pH 5.3), neutral PBS (pH 7.4) or 50 mM sodium phosphate (pH 8.3). D) Percentage of remaining active SN-38 (1.7 µM total concentration) at different incubation time points in acidic PBS (left panel) and sodium phosphate buffer (right panel) at 37 °C. E) Percentage of active SN- 38 in bulk (dashed bars), in unpurified SN-38-loaded HSPC-liposomes (grey bars) and in purified SN-38-loaded HSPC-liposomes (black bars) after dilution in basic buffer, and 0 and 2 h incubation at 37 °C. F) Schematic of the two-step liposome purification to assay SN-38 release from HSPC- liposomes. SN-38 molecules are represented as green triangles and liposomes in red. G) SN-38- containing liposomes were subjected to a two-step purification and the fraction of remaining SN-38 was calculated. The elution profiles of SN-38 (green) and liposome (red) fluorescence intensity are shown for the two chromatography steps. The elution samples that contained liposomes after the first purification were pooled and run through a second column. H) Percentage of remaining SN-38 after the second purification.

SN-38 was incubated in buffers with different pH values to populate the solution with a single or the two species. The lactone and carboxylate SN-38 forms can clearly be distinguished as two baseline-separated peaks by HPLC (Fig. 3c). The peaks eluting at 1.8–2.0 min and 6.2–6.3 min correspond to the inactive and the active forms of SN-38, respectively. At pH 5.3, approximately ∼100% of SN-38 remained in the lactone state. At pH 7.4, ∼69% of SN-38 was converted to the carboxylate form, whereas ∼95% underwent conversion at pH 8.3. In acidic PBS, SN-38 remained active for several weeks (data not shown). Maximum conversion of the lactone SN-38 to the carboxylate form in the inactivating sodium phosphate buffer (pH 8.3) was reached after 2 h (Fig. 3d).

Next, we set out to determine whether the liposomes exerted a protective effect on SN-38 against pH inactivation. We diluted free lactone SN-38, unpurified SN-38-loaded HSPC- liposomes and purified SN-38-loaded HSPC-liposomes in an inactivating basic buffer. The remaining fraction of active SN-38 present in each sample was quantified by HPLC. No significant difference was found between the three conditions after 2 h incubation (Fig. 3e), suggesting that SN-38 is released from the membrane and undergo conversion in the bulk solution. Alternatively, SN-38 interacting with, or entrapped in liposomes would get inactivated. To discriminate between these two scenarios, the SN-38 release rate from HSPC-liposomes was studied by a two-step column purification (Fig. 3f). All liposome-containing fractions in the first chromatography were pooled and subjected to a second purification after 20 min (Fig. 3g). Less than 10% of the SN-38 that had co-eluted with liposomes in the first purification step was recovered (Fig. 3h). This result shows that SN-38 is rapidly released from the HSPC liposomes into the outside solution, where it can be converted into the carboxylate form under basic conditions.

### Complexation of SN-38 with SBE-β-CD increases the drug solubility

To avoid premature release of SN-38 from HSPC vesicles, we aimed to create an SN-38-in- cyclodextrin-in-liposome system as a novel strategy for liposomal SN-38 delivery (Fig. 4a). First, we addressed the ability of SBE-β-CD to solubilize SN-38 without affecting its conversion rate. We verified that complexation with SBE-β-CD does not influence the elution profile of the lactone SN-38 by HPLC (Fig. S2). An SN-38 solubility curve was plotted by measuring the amount of solubilized drug (100 µM input concentration) as a function of SBE-β-CD concentration (Fig. 4b). The solubility of SN-38 increased linearly with respect to cyclodextrin concentration until a molar excess of SBE-β-CD to SN-38 of 1,000 was reached, after which the curved plateaued. A good agreement was found between the plate reader and HPLC methods. Maximum SN-38 solubilization is observed at a 1,000-fold molar excess of SBE-β-CD for concentrations of SN-38 ≤ 100 µM. SN-38 solubility increases by more than 70-fold compared to a molar ratio 1:8. Fluorescence-based calibration curves were constructed to enable SN-38 quantification in liposome samples from HPLC and plate reader measurements. As shown in Fig. 4c, the obtained curves were linear across four orders of magnitude of SN-38 concentrations.

**Figure 4.**
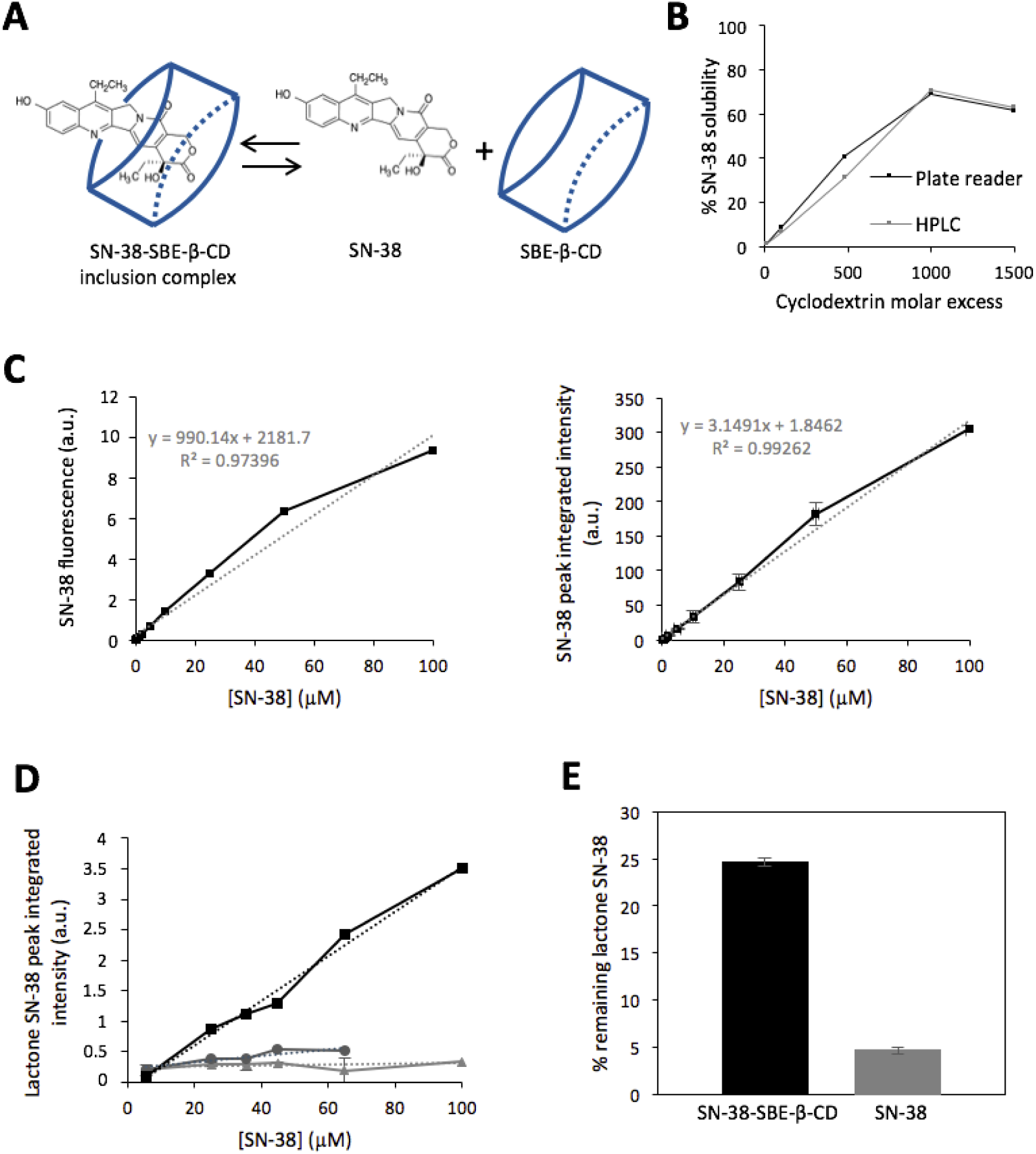
A) Schematic of SN-38 and SBE-β-CD reversible binding. B) Solubilization curves of 100 µM SN-38 by varying amounts of SBE-β-CD as measured by HPLC and on a plate reader. C) Calibration curves for quantification of SN-38 on a plate reader (left) and by HPLC (right). Eleven different SN-38 concentrations ranging from 0.01 to 100 µM were solubilized with 100 mM SBE- β-CD. The SN-38 fluorescence intensity and chromatogram peak area were measured and plotted as a function of the SN-38 concentration. The appended dotted lines represent a linear regression across the data. D) The SN-38 fluorescence peak area was measured at different SBE- β-CD concentrations by HPLC. Samples consisted of SN-38 (light grey), 1.25:1 SN-38 to SBE-β-CD molar ratio (dark grey) or a 1:8 SN-38 to SBE-β-CD ratio (black). E) Percentage of lactone SN-38 in samples with or without an excess of SBE-β-CD after 90 h incubation at room temperature in 50 mM sodium phosphate (pH 8.3). The percentage of active lactone SN-38 was calculated based on the integrated peak areas of the lactone and carboxylate species in the HPLC chromatogram. Error bars in C-E indicate mean values ± s.e.m. from three independent experiments.

The solubilization profile of SN-38 was further examined at lower relative amounts of SBE-β-CD, specifically at 1.25:1 and 1:8 SN-38 to SBE-β-CD molar ratios. The absolute concentrations of the drug and of SBE-β-CD were varied, while maintaining the ratio values constant, and the concentration of soluble SN-38 was determined by HPLC after discarding the insoluble fraction. Complexation with 64.9 µM of SBE-β-CD enhances SN-38 solubility by about 1.5 and 10 times at a 1.25:1 and 1:8 molar ratio, respectively (Fig. 4d).

We assayed whether inclusion of SN-38 in the SBE-β-CD binding cavity confers stability of the lactone ring against an inactivating surrounding buffer. The lactone SN-38 was mixed with a large excess of cyclodextrin and incubated in a basic buffer. As shown in Fig. 4e, the fraction of lactone SN-38 was almost 5 times higher after complexation with SBE-β-CD and 90 h incubation. This result demonstrates that docking of the lactone SN-38 into SBE-β-CD improves its stability, which is consistent with previous observations [28].

### Entrapment of SN-38-SBE-β-CD complexes into liposomes leads to the formation of stable SN- 38-SBE-β-CD-lipid nanoparticles

Next, we aimed to encapsulate pre-formed SN-38-SBE-β-CD inclusion complexes inside the lumen of folate-conjugated HSPC-liposomes. Because SBE-β-CD can deplete lipids [36], we anticipated that liposomes may not be stable in the presence of high concentration of either SBE-β-CD or SN-38-SBE-β-CD complexes. We first verified that the conditions to swell the lipid film did not inactivate SN-38 (Fig. S3). Then, liposomes were produced by lipid film hydration with an aqueous solution containing pre-incubated SN-38 (44 µM) and SBE-β-CD at a 1:8 molar ratio. The sample was analysed by size exclusion chromatography. As shown in Fig. 5a, the membrane dye signal was split in two distinct populations, in contrast to the single-peak profile obtained in the absence of SBE-β-CD (Fig. 3a, b). The earlier population corresponds to the typical profile of liposomes, while the later may support the presence of smaller lipid nanoparticles resulting from partly solubilized liposomes by cyclodextrin. A similar characteristic elution profile was observed when the lipid film was rehydrated with SBE-β-CD only (Fig. 5b, middle panel). We found that SN-38-SBE-β-CD inclusion complexes co-eluted with the lipid nanoparticle fractions but not with the vesicles. No changes of the co-elution profile were observed when swelling duration was extended from 2 h to overnight (Fig. S4) or when freeze-thaw cycles were applied to increase the encapsulation efficiency (Fig. 5c). Moreover, preparing lipid particles of bigger diameter (extruded across a membrane with 400-nm pores) to better accommodate SBE-β-CD-SN-38 complexes in the lumen of vesicles failed to enhance encapsulation (Fig. 5c).

**Figure 5.**
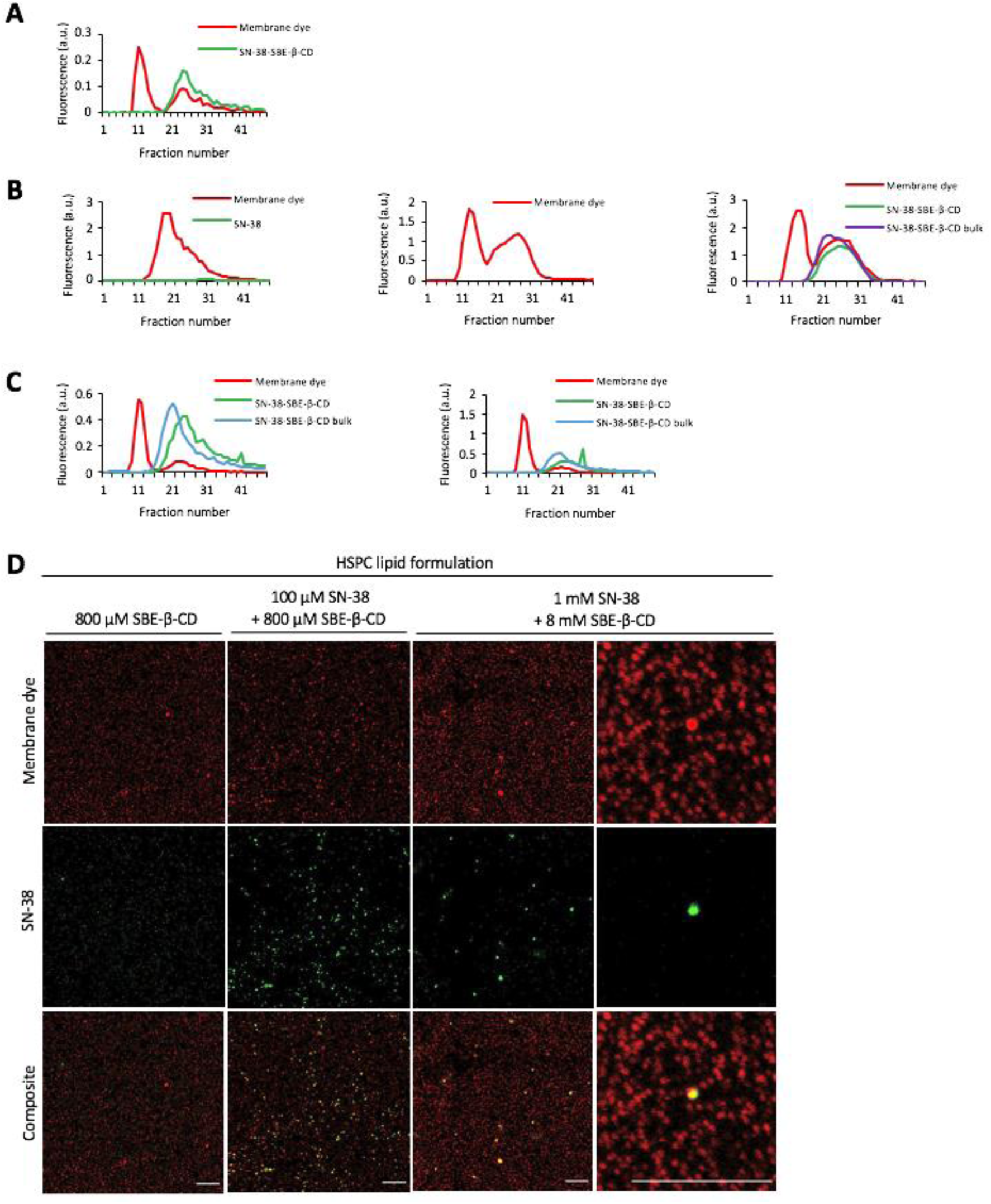
A) Elution profiles of SN-38-SBE-β-CD-loaded HSPC-liposomes after 2 h swelling (SN-38 was 44 µM, SBE-β-CD was 357.6 µM, corresponding to a 1:8 molar ratio). Fluorescence signals of Texas Red membrane dye (red) and SN-38 (green) are displayed. B) Comparison of the elution profiles with only SN-38 (left), only SBE-β-CD (middle) and with SN-38-SBE-β-CD (right). Lipid film swelling occurred for 72 h at room temperature. C) Liposomes were exposed to freeze-thaw cycles after swelling in the presence of SN-38-SBE-β-CD. Lipid particles were extruded across a 200-nm (left) or 400-nm (right) cutoff membrane. D) Confocal microscopy images of single immobilized nanoparticles from the second population of HSPC-nanoparticles entrapping 100 µM SN-38 + 800 µM SBE-β-CD, only 800 µM SBE-β-CD, or 1 mM SN-38 + 8 mM SBE-β-CD. Scale bars are 10 µm.

Based on these results, we hypothesized that the late-eluting particles are composed of water soluble lipid-cyclodextrin-SN-38 complexes as a product of lipid depletion and complexation by SBE-β-CD. We examined the elution profile of SN-38-SBE-β-CD inclusion complexes in the absence of lipids and found that it matches with the one observed in the presence of HSPC liposomes (Fig. 5b, right panel). Therefore, it is not possible to unambiguously conclude from these chromatography data that the SN-38-SBE-β-CD and lipid-SBE-β-CD inclusion complexes directly interact or assemble into mixed supramolecular structures. To further study the nature of these molecular complexes, we prepared lipid-SN-38-SBE-β-CD samples and attempted to characterize the purified fractions corresponding to the second population by fluorescence confocal microscopy. Lipid particles were immobilized on the glass surface through their biotin linker and imaged with the Texas Red dye. SN-38 fluorescence was also imaged and its localization analysed. Larger and brighter lipid particles were found to also exhibit SN-38 fluorescence, unlike the more abundant smaller and dimer structures (Fig. 5d). Increasing the concentration of SBE-β-CD from 800 µM to 8 mM (the concentration of SN-38 was also increased 10 times to keep the same molar ratio) did not affect the apparent size of the lipid aggregates (Fig. 5d). These results of visualization of single particles suggest direct interaction between SN- 38-SBE-β-CD and lipid-SBE-β-CD inclusion complexes, at least for the bigger structures. The fact that these lipid-SN-38-SBE-β-CD aggregates retained a fraction of SN-38 even 24 h after dilution in physiological buffer (PBS pH 7.4) through size exclusion chromatography prompted us to explore their antitumor effect *in vitro*.

### SN-38-SBE-β-CD-lipid nanoparticles can interact with cancer cells

The first step in the antitumor mechanism of a therapeutic lipid nanoparticle is the interaction with the plasma membrane and subsequent internalization. Thus, we investigated the interaction of SN-38-SBE-β-CD lipid nanoparticles with cultured KB cells. Purified lipid nanoparticles from the second population obtained by size exclusion chromatography were incubated with plated living cells in a growth medium for up to 2 h. Though the yield of nanoparticles was very low due to the loss of material during column purification, the interaction of a few SN-38-SBE-β-CD-lipid nanoparticles with KB cells was observed by time-lapse confocal microscopy (Fig. 6a). The fluorescence signal of SN-38 was too low to be measured in cells.

**Figure 6.**
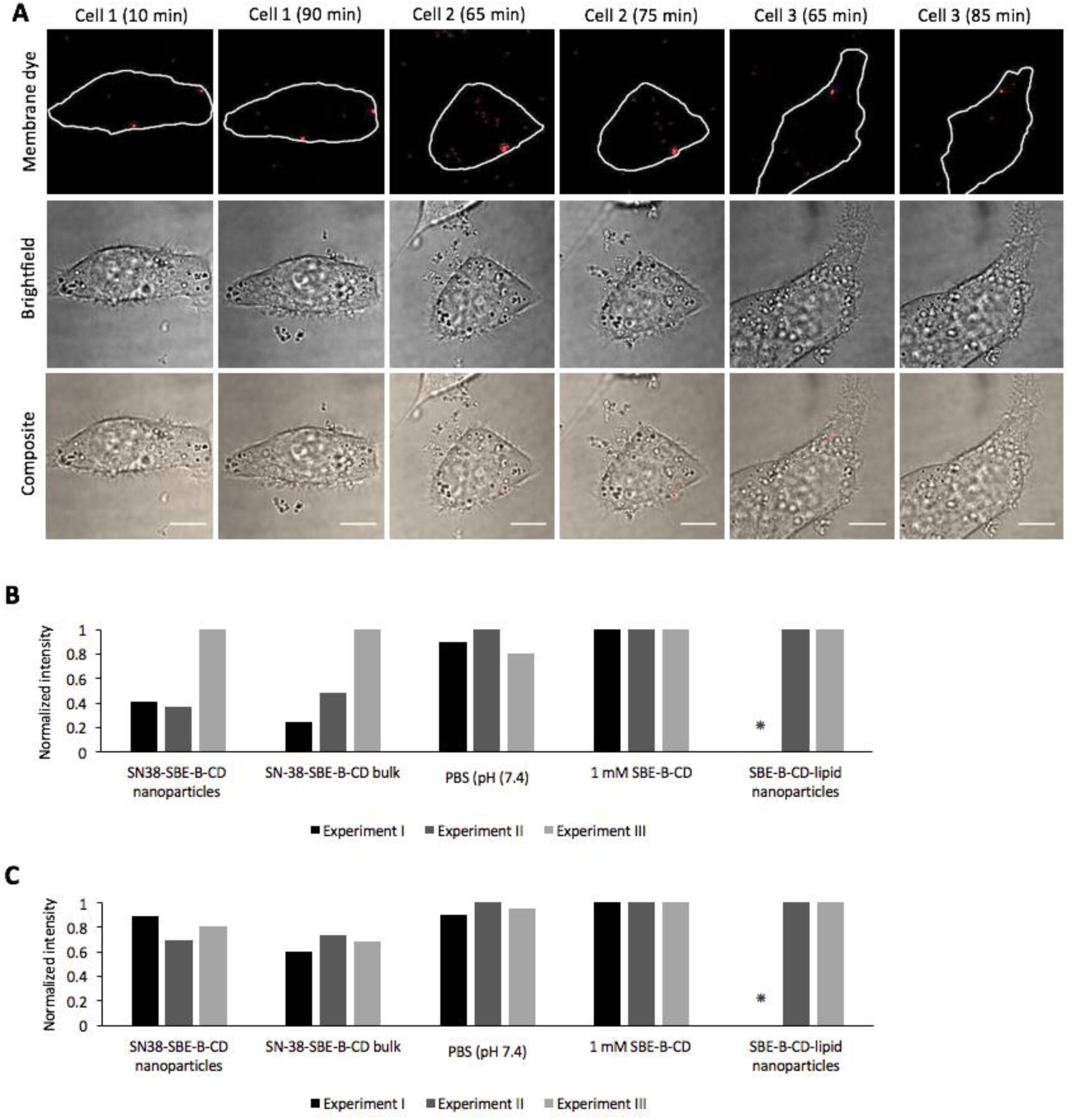
A) Confocal images of SN-38-SBE-β-CD-lipid nanoparticles interacting with KB cells at different time points. Scale bars are 10 µm. B,C) Normalized cell viability of B) KB and C) HeLa cells after 72 h incubation with either SN-38-SBE-β-CD-nanoparticles, bulk SN-38-SBE-β-CD, SBE- β-CD-lipid nanoparticles, 1 mM SBE-β-CD or PBS pH 7.4. The asterisk indicates that this experimental condition was not conducted. Three independent experiments were performed.

To evaluate the *in vitro* toxicity of SN-38 as an SN-38-SBE-β-CD-lipid nanoparticle formulation, the purified particles were incubated for 24 h with KB and HeLa cells. Control experiments were performed with SN-38-SBE-β-CD inclusion complexes. Fig. 6b and 6c show the results from three independent experiments. The toxicity effects on KB cells in the third experiment significantly differed from the other two. Nevertheless, the antitumor effect of SN-38-SBE-β-CD-lipid nanoparticles was similar to that of the SN-38-SBE-β-CD complexes in both cell types for all experiments. Importantly, SBE-β-CD-lipid nanoparticles (no SN-38) did not induce cytotoxicity, strongly suggesting that this range of cyclodextrin concentrations is safe for drug delivery and will not lead to unspecific effects. In addition, HeLa cells were on average less sensitive to the effect of SN-38 than KB cells. Altogether, we attribute the lack of toxicity to the low concentration of SN-38-SBE-β-CD-lipid nanoparticles used in these experiments, a consequence of the large dilution during size exclusion chromatography.

## DISCUSSION

In spite of its activity against tumor cells *in vitro*, the clinical use of SN-38 (and other hydrophobic compounds) for chemotherapy remains limited due to its limited solubility and instability at physiological pH. To solve these problems, a variety of liposomal formulations have been developed [6]. In recent years, drug-in-cyclodextrin-in-liposomes have been designed to further improve the delivery properties [33]. Complexation of SN-38 with SBE-β-CD greatly increases its solubility [28]. In the present study, the development of SN-38-in-cyclodextrin-in-liposomes has been explored. We have attempted to equip this system with folic acid as a ligand for targeted delivery and with polyethylene glycol to manage highly stable, long-circulating lipid particles.

With the ultimate goal of developing liposomes that selectively deliver SN-38 to tumor cells, we developed a formulation of folic acid-labelled liposomes. For more than two decades, folate receptors have been intensively studied as tumor-specific drug delivery targets [37–40]. Here, we demonstrated selective uptake of folate-labelled HSPC-liposomes by cultured cells expressing folate receptors (Fig. 2c). *In vivo*, this would translate into an increased tumor accumulation of liposomes compared to non-targeted liposomes. However, the degree of increase is highly variable and *in vivo* studies would be required to evaluate the tumor targeting potential of this liposome formulation to folate receptor-overexpressing tumors [40].

As a first approach to solubilize and protect SN-38, we developed a formulation of SN-38-loaded- HSPC liposomes. The production of lipid-coated beads enabled the optimal storage of the SN- 38-lipid formulation for months and presumably leads to a higher concentration of generated liposomes and an enhanced drug-encapsulation efficiency thanks to the increased surface area provided by the glass beads compared to ordinary methods [41, 42]. Preliminary liposome characterization by fluorescence microscopy and dynamic light scattering demonstrated that the presence of SN-38 did not interfere with the formation or stability of these lipid vesicles (Fig. 2a, b). This was an important verification considering that some lipophilic drugs are known to interfere with bilayer formation and stability upon entrapment [23, 24]. The observed variability on the fractions at which liposomes eluted during size exclusion chromatography can be attributed to inherent differences between the purification columns (e.g. resine packing, amount of resine), as they were handmade for each sample (Fig. 3a, b). Successful entrapment of some SN-38 molecules in HSPC-liposomes was demonstrated and a larger fraction of SN-38 was shown to interact if the sample was incubated prior to purification (Fig. 3b). Since the buffer used to generate the liposomes was acidic, all SN-38 molecules were in the closed lactone ring conformation. Due to the high hydrophobicity of the lactone SN-38, the drug is expected to be buried in the lipid bilayer [3]. In particular, the long 18-carbon acyl chains of HSPC could favor the entrapment of SN-38 into the membrane. A different formulation of SN-38-loaded liposomes, named LE-SN-38, has previously been created and SN-38 was found to localize in the outer leaflet of the liposome bilayer, at the water interface [3]. In our study, the rapid release of SN-38 from HSPC liposomes indicates a weak interaction (Fig. 3e). Moreover, the non-significant difference in the fraction of active SN-38 between purified liposomes, non-purified liposomes and bulk SN-38, all incubated in an inactivating alkaline buffer, further indicates that HSPC-liposomes do not protect SN-38 from external protons. Hence, SN-38 might not be stably buried in the core of the lipid bilayer, but may instead interact near the liposome surface, as shown for LE-SN-38.

To circumvent the short retention time of SN-38 by liposomes, we aimed to develop a drug-in- cyclodextrin-in-liposome system that would encapsulate drug-cyclodextrin complexes within the lumen of the vesicle. Consistent with previous results [28], the solubility and stability of SN- 38 increased upon formation of inclusion complexes with SBE-β-CD (Fig. 4). Current marketed Captisol-enabled drug formulations have molar cyclodextrin excesses between 1.4 and 4.3. Cyclodextrins have the ability to interact with and extract cholesterol [43] and other membrane lipids, altering the properties of cellular membranes and inducing cytotoxic effects [36]. Mindful that large amounts of cyclodextrin could be lethal to cells, we first characterized the solubilization of SN-38 at low SBE-β-CD to drug molar ratios, and found that a value of 1:8 offered a good comprise between effective solubilization of SN-38 and absolute cyclodextrin concentration for further cytotoxicity assays (Fig. 4c).

Though the concept of drug-in-cyclodextrin-in-liposomes as drug delivery system was first proposed more than two decades ago [24], cyclodextrin-drug complexation competes with liposomal membrane assembly, tempering the potential benefit of complexation in prolonging hydrophobic drug retention [44]. Earlier studies on the effect of different cyclodextrins on the integrity of liposomes reported that vesicles can collapse due to cyclodextrin-induced extraction of lipids, causing the formation of water soluble lipid-cyclodextrin complexes. Moreover, this effect depends on the cyclodextrin type and the length of the lipid fatty acid chain [43, 45, 46]. Herein, we showed that SN-38-in-SBE-β-CD-in-liposomes fail to self-assemble at 1:8 drug to cyclodextrin molar ratio (Fig. 5). Instead, size exclusion chromatography and fluorescence confocal microscopy revealed that SN-38 is trapped in lipid nanoparticles, likely arising from a cyclodextrin-induced lipid extraction mechanism. Under certain conditions, cyclodextrin molecules and their ligands can form non-inclusion aggregates with a size ranging from 20 to 100 nm in diameter [47]. We hypothesized that the observed nano-aggregates comprise SN-38- SBE-β-CD and lipid-SBE-β-CD inclusion complexes. Their nearly identical elution profiles suggest that both types of particles have similar sizes (Fig. 5c). It is known that the aggregation propensity and the size of the aggregates increase with increasing cyclodextrin concentration [47]. The optical microscopy experiments performed here should be complemented with transmission electron microscopy measurements to determine the size, morphology and exact molecular structure of these nanoparticles.

Liposomes are intrinsically highly heterogeneous assemblies, exhibiting different physical and chemical properties (e.g. size, lipid composition, encapsulation efficiency) even within a same batch [48]. Therefore, successful implementation of liposome-based drug delivery systems requires characterization of those properties at the single nanoparticle level. Transmission electron microscopy is a powerful technique to characterize the size, morphology and lamellarity of individual liposomes [48] and can be used as well for examining the heterogeneity of drug encapsulation yield in vesicles [49, 50]. However, this technique necessitates laborious sample preparation and allows the analysis of only a small fraction of the sample. Fluorescence microscopy of immobilized liposomes or nanoparticles does not provide accurate information about the size or morphology of molecular assemblies with dimensions lower than about 200 nm. However, it enables high-throughput evaluation of the drug loading heterogeneity between individual liposomes provided the drug has fluorescent properties, as demonstrated in this study for SN-38 (Fig. 5d). The use of fluorescence microscopy of surface-immobilized vesicles in the field of liposomal drug delivery development has already been proposed [48, 51], but had not been applied yet.

Due to the low concentration of purified SN-38-SBE-β-CD-lipid nanoparticles, we were unable to reliably assay their interaction dynamics with cultured KB cells and to visualize the fate of SN-38 after binding to the plasma membrane. Although we primarily attribute the lack of enhanced cytoxicity of SN-38-SBE-β-CD-lipid nanoparticles compared to non lipidic formulations of SN-38 to their low concentration, we cannot exclude that other factors may contribute too, such as poor nanoparticle uptake due to low folic acid ligand availability on their surface, poor endosomal escape or release of SN-38 in the cytoplasm, and SN-38 inactivation.

## CONCLUSIONS

Together, our results provide clear directions for further nanoparticle characterization and physico-chemical optimization of the SN-38 formulation. Electron microscopy will be an essential tool to understand the molecular nature and size of the observed nano-aggregates. Furthermore, increasing the amount of HSPC-liposomes relative to that of SN-38-SBE-β-CD complexes might lessen the lipid bilayer depletion effect and facilitate the successful production of drug-in-cyclodextrin-in-liposomes. Alternatively, the lipid-extraction effect of other SN-38 solubilizing β-cyclodextrins such as randomly methylated β-cyclodextrin (RMβCD, Trappasol) or methylated β-cyclodextrin (MβCD) [28] should be explored. Finally, increasing the concentration of purified SN-38-SBE-β-CD-lipid nanoparticles, currently a bottleneck for cellular cytotoxicity, might be achieved by scaling up the production and by concentrating the particles by ultracentrifugation or solvent evaporation.

## Acknowledgements

We thank Esengül Yildirim for assistance with HPLC experiments, Esther Hulleman, Dennis Metselaar and Lot Sewing for discussions about SN-38. This work was financially supported by the FOM programme nr.140.

## Author Contributions

AS and CD designed experiments, analysed data and wrote the manuscript. AS performed experiments. NB contributed methods development to detect SN-38 by HPLC.

## Supplementary Information

Supplementary Figures are available.

