## Supplementary Information for "Development of a lipid-based delivery system for the chemotherapeutic compound SN-38"

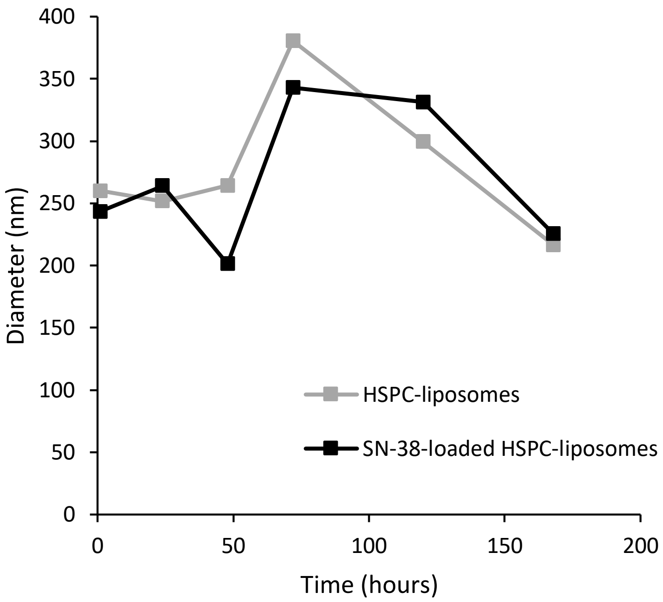


**Fig. S1.** Dynamic light scattering data showing the liposome size distribution of SN-38-loaded HSPC-liposomes and of HSPC-liposomes at different storage time points.

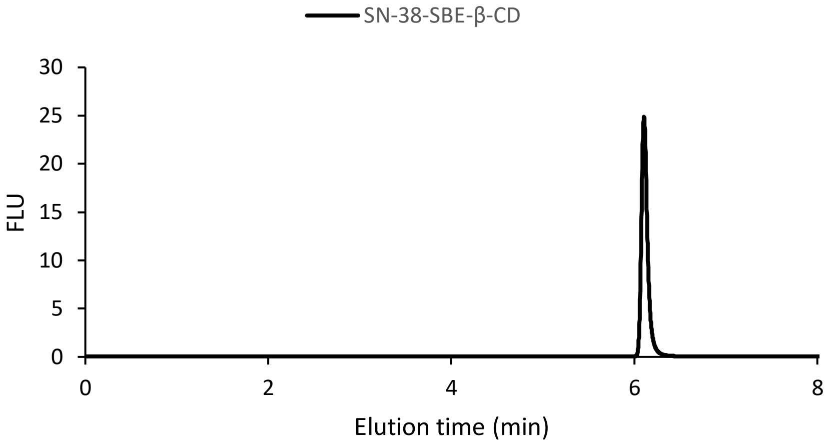


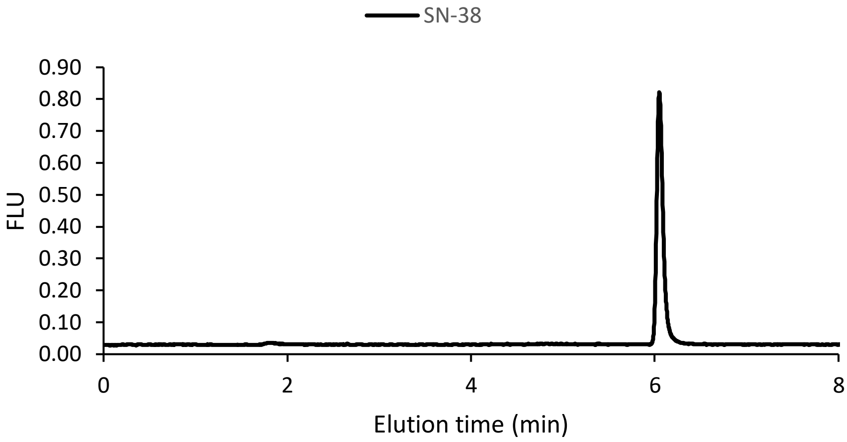


**Fig. S2.** HPLC chromatograms of SN-38-SBE-β-CD (top) and SN-38 (bottom).

To discard a possible influence of SBE-β-CD complexation on the elution profile of SN-38, we incubated SN-38 (25 µM) at room temperature overnight in the presence or absence of a large excess of SBE-β-CD (4.8 mM, 1:192 molar ratio). The resulting elution time and shape of the lactone SN-38 peak were analyzed by HPLC. As expected, due to the non-covalent interaction between SN-38 and SBE-β-CD, the elution time of lactone SN-38 remained the same in both samples.

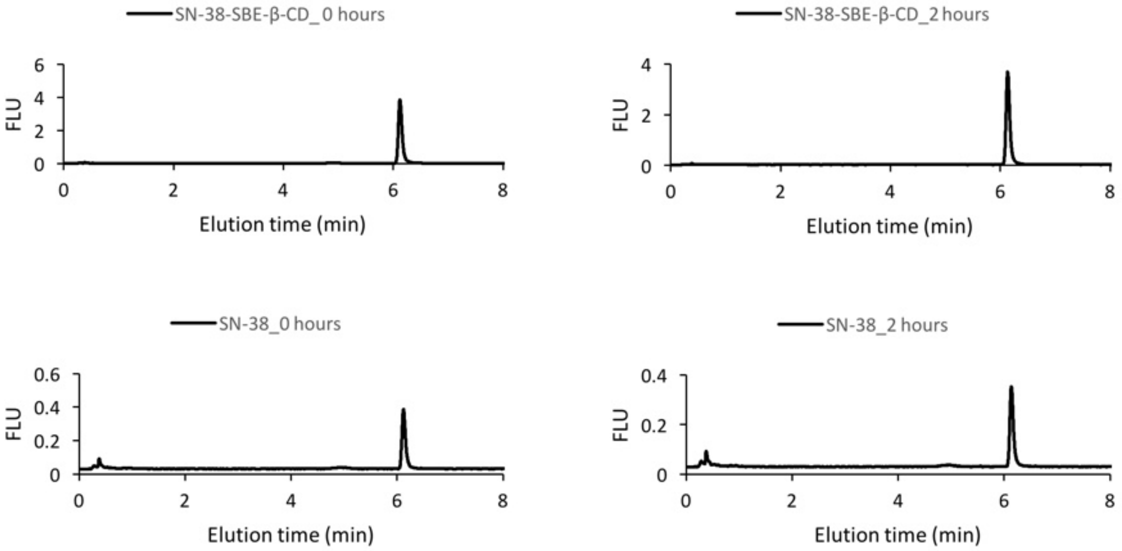


**Fig. S3.** Stability of SN-38 at 60 °C in the presence or absence of SBE-β-CD. HPLC chromatograms of SN-38-SBE-β-CD and SN-38 without and with 2 h incubation at 60 °C. Concentrations of SN-38 and SBE-β-CD in solution were 100 µM and 800 µM, respectively.

A solution containing 100 µM SN-38 plus 800 µM SBE-β-CD and a solution containing 100 µM SN-38 were prepared in acidic PBS (pH 5.3). The tubes were sonicated for 15 min, incubated under tumbling at room temperature overnight and passed through a 0.45-µm filter. The tubes were then sealed and incubated at 60 °C for 2 h in a shaker at 250 rpm. 10 µl were injected into the HPLC and the amounts of the lactone and carboxylate forms of SN-38 were measured. Complexed and non-complexed SN-38 remained in the lactone form after 2 h incubation at 60 °C. Moreover, the shape of lactone SN-38 peaks after incubation was as sharp as at time point 0, confirming that this condition does not harm the structure of SN-38. Consequently, swelling for up to 2 h at temperatures equal or lower than 60 °C does not affect the structure of SN-38.

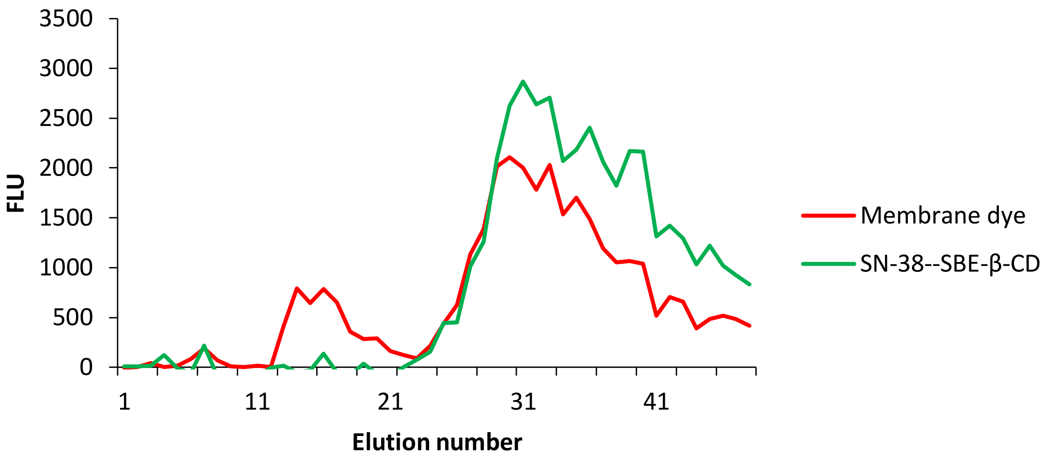


**Fig. S4.** Elution profiles of SN-38-SBE-β-CD-loaded HSPC-liposomes after overnight swelling (SN-38 44 µM, SBE-β-CD 357.6 µM, 1:8 molar ratio).
